# Dead or Alive? Molecular life-dead distinction in human stool samples reveals significantly different composition of the microbial community

**DOI:** 10.1101/343194

**Authors:** Alexandra Perras, Kaisa Koskinen, Maximilian Mora, Michael Beck, Lisa Wink, Christine Moissl-Eichinger

## Abstract

The gut microbiome is strongly interwoven with human health. Conventional gut microbiome analysis generally involves 16S rRNA gene targeting next generation sequencing (NGS) of stool microbial communities, and correlation of results with clinical parameters. However, some microorganisms may not be alive at the time of sampling, and thus their impact on the human health is potentially less significant. As conventional NGS methods do not differentiate between viable and dead microbial components, retrieved results provide only limited information.

Propidium monoazide (PMA) is frequently used in food safety monitoring and other disciplines to discriminate living from dead cells. PMA binds to free DNA and masks it for subsequent procedures. In this article we show the impact of PMA on the results of 16S rRNA gene-targeting NGS from human stool samples and validate the optimal applicable concentration to achieve a reliable detection of the living microbial communities.

Fresh stool samples were treated with a concentration series of zero to 300 μM PMA, and were subsequently subjected to amplicon-based NGS. The results indicate that a substantial proportion of the human microbial community is not intact at the time of sampling. PMA treatment significantly reduced the diversity and richness of the sample depending on the concentration and impacted the relative abundance of certain important microorganisms (e.g. *Akkermansia, Bacteroides*). Overall, we found that a concentration of 100 μM PMA was sufficient to quench signals from disrupted microbial cells.

The optimized protocol proposed here can be easily implemented in classical microbiome analyses, and helps to retrieve an improved and less blurry picture of the microbial community composition by excluding signals from background DNA.

## Introduction

The function of microbial communities depends on which community members are dead or alive, and the information about their status can be crucial for questions concerning microbial ecology or clinical relevance. For example a viable pathogenic bacterium poses a different risk to humans when present in pharmaceuticals or food as it might cause infection ^1^, whereas contamination with dead microorganisms poses a potential risk for intoxication. Furthermore, clinical efficiency of fecal microbiota transplants depends on the living bacterial fraction of the preparation as only these bacteria could colonize the gut of the receiving patient ^2^.

Several techniques have been developed to differentiate between live and dead microbes, focusing on three basic principles: To be considered alive, a microbial cell must be (1) intact, (2) able to reproduce, and (3) metabolically active (Emerson et al, 2017). The capability to reproduce can be ultimately assessed by cultivation-dependent methods, as, for the formation of colony forming units, cells have to divide and grow. However, only a comparatively small fraction of the Earth’s microbiology can be cultivated using standard laboratory techniques at all ^3, 4^. Metabolic activity can be estimated by metabolomics analyses or by analysing the content of (r)RNA, as it reflects the activity of microbial transcription; however, also dead cells can contain a considerable amount of RNA, blurring the results and conclusions ^5^. Besides these vague approaches, the intactness of a cell is an unquestionable prerequisite for life, as life outside of a cell compartment does not exist.

One methodology to assess the intactness of a cell relies on the application of propidium monoazide (PMA). PMA masks free DNA and DNA of non-intact cells ^6^, as it can pass disrupted cell walls and intercalate into the DNA double helix. Upon light exposure, PMA is covalently bound, blocking this DNA from subsequent PCR amplification ^6^. When applied prior to DNA extraction, sequencing or qPCR, PMA treatment results in exclusive detection of the intact microbial fraction. By this principle, PMA treatment may also improve detection of rare community members as it helps to exclude background DNA from analysis ^7^. PMA has been applied in a wide variety of environmental and clinical microbiome studies, including research on the human microbiome ^7-9^, indoor environments and built systems ^10, 11^ as well as on food products ^12-14^. PMA is most typically used in standard PCR-based assays, but has also been successfully applied in combination with e.g. qPCR^15-17^, metagenomics^18^, fluorescence *in situ* hybridization (FISH)^19^, fluorescence microscopy^20^ and microarrays ^21^. Another application aimed at the treatment of commercial PCR reagents to mask DNA contaminations (the “kitome”;^12^.

The original method foresees sample treatment with defined PMA concentration (50 μM), incubation in the dark (5 min, agitation) and a subsequent light exposure (2 min, 650 W lamp;^22^. However, the efficiency of PMA treatment is highly affected by density, homogeneity and turbidity of the handled sample ^23, 24^ and thus the original methodology might not be applicable to e.g. stool samples, although PMA treatment of faeces samples is as well a desired goal and has already been applied ^2, 25, 26^. However, despite the high biomass and the complexity of faeces, optimal conditions of PMA treatment of faeces samples have not been elaborated yet. As a consequence, we wanted to know, whether an increased PMA concentration could compensate the problems of turbid and complex biosamples, such as faeces.

We herein analyze the effect of increasing PMA concentrations on the microbial community composition in human stool samples using amplicon-based next generation, with the goal to establish a guideline on how to reliably apply PMA in complex biosamples.

## Material and Methods

### Samples and general organisation of experiments

Fresh human feces was obtained from a healthy human individual and kept on ice until processing. The samples were diluted 1:10 (w/v) in DNA-free, sterile 1x phosphate buffered saline (PBS), and 500 μl aliquots of the thoroughly mixed dilution were treated separately with increasing concentrations of PMA. We performed two subsequent experiments, experiment A and B. In experiment A, we studied the effects of PMA with 11 different concentrations between 0 μM and 100 μM PMA, and in experiment B we tested five different concentrations of PMA between 0 μM and 300 μM. The fecal samples for experiment A and B were obtained from the same person, but at different time-points. The samples were sequenced in the same Illumina MiSeq run.

### PMA cross-linking and DNA isolation

PMA was dissolved in PCR grade water to create a stock concentration of 20 mM and stored at −20°C in dark until usage. PMA was added directly to 500 μl sample aliquots applying the following PMA concentrations in experiment A: 0 μM, 10 μM, 20 μM, 30 μM, 40 μM, 50 μM, 60 μM, 70 μM, 80 μM, 90 μM and 100 μM, and the following concentrations in experiment B: 0 μM, 50 μM, 100 μM, 150 μM, and 300 μM. The samples were shaken on ice in dark for 10 minutes (IKA Rocker, 60 rpm) and light-exposed for 15 minutes in a standard PMA-Lite™ LED photolysis device with occasional shaking. After photo-induced cross-linking, DNA was extracted using the PowerSoil DNA extraction kit (Mo Bio Lab, Inc.) following the manufacturer’s instructions. DNA concentrations were measured using the Qubit (Thermo Fisher Scientific) method and each assay was diluted to 10ng/μl in PCR grade water as template for subsequent PCR reactions. All experiments were performed in triplicates. DNA extraction blank samples were used to control for the sterility of reagents and equipment.

### Library preparation for next generation sequencing (NGS)

Variable region V4 of the 16S rRNA gene was amplified with universal PCR primers 515F (5’-GTGCCAGCMGCCGCGGTAA-3’) and 806R (5’-GGACTACHVGGGTWTCTAAT-3’) using TaKaRa Ex Taq polymerase (Clontech, Japan) ^27^. The cycling conditions were: initial denaturation at 94 °C for 3 min, followed by 35 cycles of denaturing at 94 °C for 45 s, annealing at 60 °C for 60 s, and elongation at 72 °C for 90 s. The final elongation step lasted 10 min. Negative controls (no template) were included in PCR. Library preparation and sequencing were carried out at the Core Facility Molecular Biology at the Center for Medical Research, Graz, Austria. Briefly, DNA concentrations were normalized using a SequalPrep™ normalization plate (Invitrogen), and each sample was indexed with a unique barcode sequence (8 cycles index PCR). Pooled samples were purified with gel-cut procedure. Sequencing was conducted using Illumina MiSeq device. Sequence data were deposited in The European Nucleotide Archive (ENA) with study accession number PRJEB25855.

### Sequence data processing and analysis

Raw sequence reads were preprocessed and filtered using the R package dada2 (version 1.4.0) according to the proposed processing pipeline ^28^. Briefly, reads were demultiplexed, forward and reverse reads were quality filtered (min. score: 30), merged, and the dada2 core algorithm was applied. Taxonomic affiliations were assigned according to SILVA v123 database^29^. All ribosomal sequence variants (RSVs) which were present in negative controls were removed from the dataset; Normalization was performed by transformation to relative abundances. The downstream analyses were performed using R software package (version 1.0.136)^30^. Alpha and beta diversities were calculated using the R packages phyloseq (version 1.20.0) ^31^ and vegan (version 2.4)^32^.

To compare microbial diversity in samples treated with different PMA concentrations, alpha diversity was calculated using the Shannon, Observed and Inverse Simpson index. Normal distribution was tested by the Shapiro-Wilk test, and statistical significance was tested either by Kruskal-Wallis (if not normally distributed) or ANOVA (if normally distributed). The Dunn’s test was used as post-hoc test (Benjamin Hochberg correction).

Beta diversity analyses were performed to examine differences between microbial community profiles at different PMA concentrations. Principal coordinate analysis (PCoA) was performed to study the relationships and variation between samples. Statistical significance was tested using adonis (seed 1) and further confirmed by dispersion test (permutations: 999). Nonmetric multidimensional scaling (NMDS) was applied to visualise these differences. The data were also subjected to linear discriminant analysis effect size (LEfSe) analysis ^33^ to determine specific microbial markers for PMA treatment of different concentrations.

## Results

In order to determine the influence of increasing PMA concentration on the microbial community profile and diversity, we carried out two experiments: In experiment A, we compared the concentrations of 10 μM to 100 μM PMA (in 10 μM steps) to non PMA treated samples and analysed the possible effects of PMA, and in experiment B, we tested the concentrations of 50 μM, 100 μM, 150 μM and 300 μM PMA to confirm experiment A. In both experiments we compared the PMA treated samples to non-treated samples.

Obtained sequence data were processed using dada2^28^, and 1,597,041 sequence reads were obtained for experiment A (33 samples plus controls), and 716,611 sequence reads for experiment B (15 samples plus controls). After removing all taxa which were present in negative controls, 1,596,911 RSV counts (unique sequence reads) affiliated to 437 different taxa were left for experiment A, and 715,808 RSV counts (unique sequence reads) affiliated to 329 different taxa for experiment B. These data were analysed to show how the PMA treatment with different concentrations can influence alpha and beta diversities and the microbial community profile in specific taxonomic groups.

The analysis revealed a typical human gut microbial community profile within the studied samples. In experiment A we identified 16 microbial phyla, of which 13 were affiliated to Bacteria and 3 were affiliated to Archaea (Euryarchaeota, MSC (miscellaneous Crenarchaeotic Groups), and Thaumarchaeota). The most abundant phyla were Bacteroidetes (52.2 % of all sequence reads), Firmicutes (37.7%), and Proteobacteria (4.4%). At genus level, the most abundant identified taxa were *Bacteroides* (37.7% of all sequence reads), *Faecalibacterium* (6%), and *Akkermansia* (4%).

In experiment B, we identified 11 microbial phyla of which 9 were affiliated to Bacteria and 2 to Archaea (Euryarchaeaota and Thaumarchaeota). Again, the most abundant phyla were affiliated to Bacteroidetes (64.3% of all sequence reads), Firmicutes (29.1%), and Proteobacteria (5.2%), while the most abundant genera were *Bacteroides* (42.9%), a taxon assigned to *Bacteroidales* family *S24* (10.1%), and *Alistipes* (5.4%). The most abundant phyla identified in experiment A and experiment B are depicted in Figure 1.

**Figure 1.**
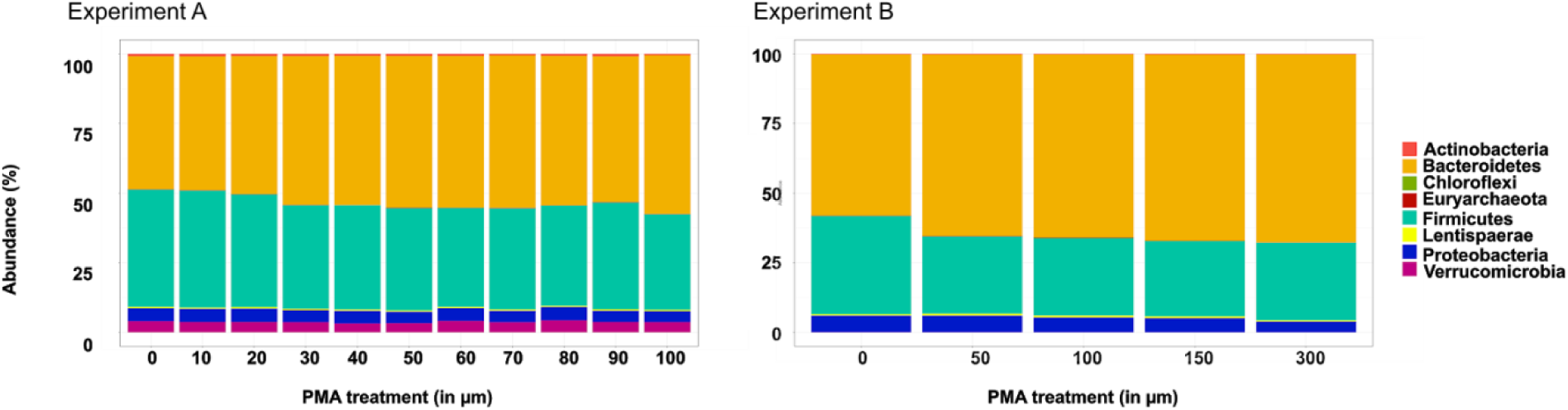
The most abundant phyla in experiment A and B. Majority of sequence reads were affiliated to Bacteroidetes and Firmicutes. Low abundant phyla (>0.1%) are not included in the figure.

### PMA treatment significantly decreases microbial richness and diversity (alpha diversity)

We studied how the PMA treatment might affect the alpha diversity of microbial communities in stool samples. The results show that alpha diversity (observed richness, Shannon and inverse Simpson indices) overall decreased with PMA treatment and that the effect was PMA concentration dependent. In experiment A the observed richness significantly decreased (ANOVA, p=1.57e-06) with 30 μM, 40 μM, and 50 μM PMA concentrations when compared to non PMA treated control (all p-values were <0.05, Tukey’s post-hoc test, confidence level 0.95). Additionally, the Shannon diversity index decreased with 100 μM PMA treatment compared to untreated samples (ANOVA, p=0.00767).

In experiment B, a significant decrease in diversity was detected in the Shannon index when 300 μM PMA was used (p=0.03121), and with inverse Simpson indices when 150 μM PMA (p=0.0498812) and 300 μM PMA (p=0.0117023) was used, compared to 0 μM PMA samples. Figure 2 summarizes the results, and highlights all statistically significant changes in alpha diversity upon PMA treatment.

**Figure 2.**
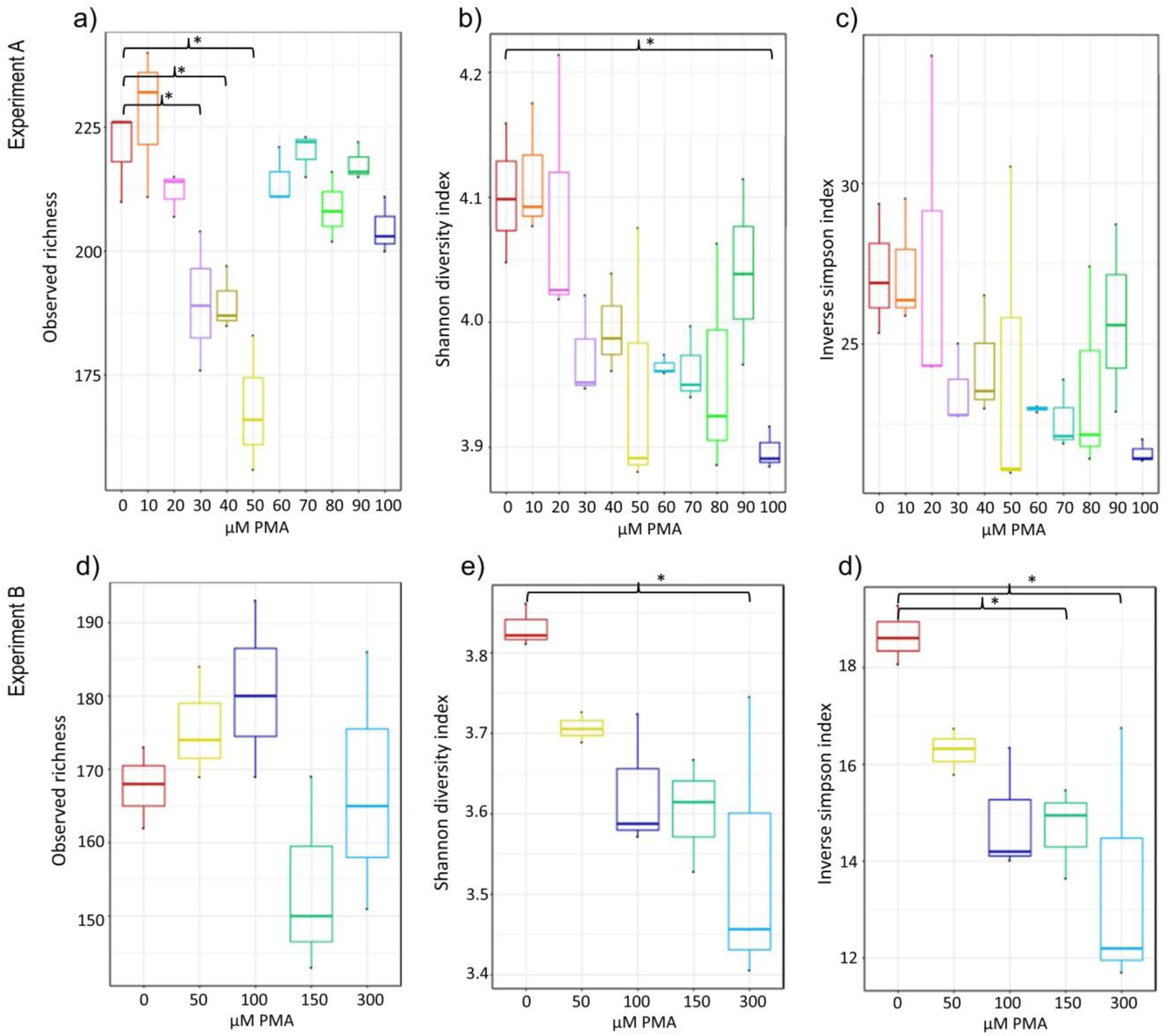
PMA treatment significantly decreases the detectable microbial diversity. In experiment A the observed richness (a) and Shannon diversity (b) significantly decreased with PMA treatment. In experiment B a significant decrease in diversity was detected by Shannon (e) and inverse Simpson (f) indices. Experiment A is pictured in the upper panel (a, b and c), and experiment B in the lower panel (d, e f). Statistically significant changes are marked with *.

### PMA treatment significantly alters the microbial community profile (beta diversity)

We studied the effect of PMA treatment also on the overall microbial community profile. Therefore, we applied various beta diversity measures, including Bray-Curtis distance (unweighted) and tested for significant differences using Adonis, confirmed with a dispersion test. We visualised the beta community profile via nonmetric multidimensional scaling (NMDS).

The effect of PMA treatment was significant with respect to concentration. The results show that PMA treatment, particularly in higher concentrations, significantly shifted the microbial community profile (Figure 3, PCoA plots) (Adonis p<0.001), from 0 μM PMA to higher concentrations (Figure 3, NMDS). The significance was emphasized with a dispersion test (permutations: 999; both p-values >0.8).

**Figure 3.**
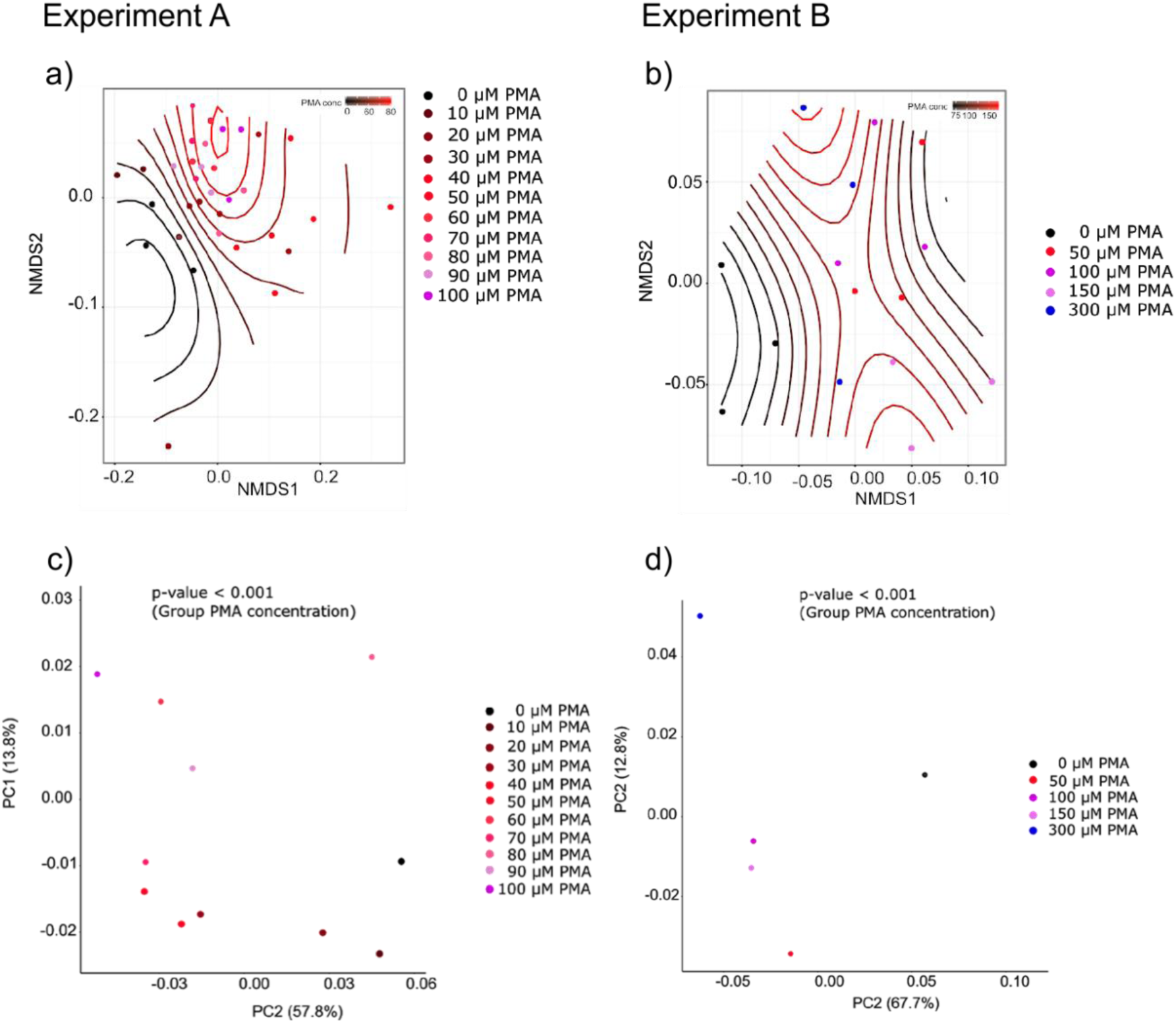
PMA treatment significantly alters the microbial community profile. Non-metric multidimensional scaling (NMDS) (a, b) visualisation presents a clear gradient from 0 μM PMA to higher concentrations. Principal coordinate analyses (PCoA) (c,d) show that PMA treatment, particularly in higher concentrations, significantly shifted the microbial community profile (Adonis p<0.001.) Experiment A is pictured in the left panel (a, c), and experiment B in the right panel (b, e).

Additionally, we were interested in which taxa are responsible for the community shift. Using linear discriminant analysis effect size (LEfSe) algorithm we compared the median values of 100 most abundant microbial RSVs between samples without PMA treatment and different PMA concentrations. We were interested in particular if certain taxa are significantly less abundant in PMA treated samples, suggesting they are in non-intact (i.e. dead) state and consequently often overestimated regarding their relative abundance in microbiome studies, or if particular taxa are not affected by the treatment, or increase in relative abundance, indicating they are present in stool samples as intact, viable cells.

In experiment A, the 100 most abundant RSVs represented 7 phyla, and in experiment B 6 phyla, which were the same as in experiment A except the phylum Verrucomicrobia, represented by one RSV of *Akkermansia.*

PMA treatment did not completely remove any taxon or RSV in experiment A, but in experiment B a total of 5 RSVs were not detected after PMA treatment, indicating these taxa were in nonviable state (Supplementary Table 2). In both experiments, the majority of the 100 most abundant RSVs either increased or decreased statistically significant (p<0.05) in relative abundance at one or more PMA concentrations (experiment A 75% RSVs changed significantly in their relative abundance, in experiment B 68%).

All RSVs that changed significantly in experiment A or B are presented in Table 1, and information including all identified phyla and genera in both experiments can be found in Supplementary table 1 and Supplementary table 2.

In this study we show that the effect of PMA treatment might be indeed taxon specific (Table 1, Supplementary table 1, Supplementary table 2). The majority (57% in experiment A / 53% in experiment B) of RSVs belonging to the phylum Firmicutes had a lower relative abundance after PMA treatment (Supplementary table 1, Supplementary table 2). In both experiments, all RSVs of the genus *Faecalibacterium,* and most RSVs in the families Lachnospiraceae and Ruminococcaceae were significantly reduced by PMA treatment, suggesting their relative abundance can be overestimated when nonviable cells are not masked (Table 1).

In contrast to Firmicutes, relative abundance of the phylum Bacteroidetes increased (69% of RSVs in experiment A / 68% of RSVs in experiment B). The majority of RSVs belonging to the genera *Alistipes* (75% in experiment A / 100% in experiment B), *Bacteroides* (71% in experiment A / 77% in experiment B), and *Parabacteroides* (50% in experiment A / 100% in experiment B) show significant increase in relative abundance after PMA treatment (Table 1, Supplementary table 1, Supplementary table 2). Additionally, the phylum Lentisphaerae (genus *Victivallis*) increased significantly in relative abundance in both experiments.

Change in relative abundance of *Bifidobacterium* (Actinobacteria) and *Akkermansia* (Verrucomicrobia) upon PMA treatment was only detectable in experiment A as these taxa were not found in experiment B. The relative abundance of *Bifidobacterium* decreased significantly when the sample was treated with 60 μM PMA or more; the relative abundance of *Akkermansia* it reduced significantly when the sample was subjected to at least 30 μM PMA treatment.

The results suggest that if PMA treatment is not applied, the relative abundance or relevance of these bacteria, which are known to carry beneficial properties for human health, could be overestimated.

**Table 1.**
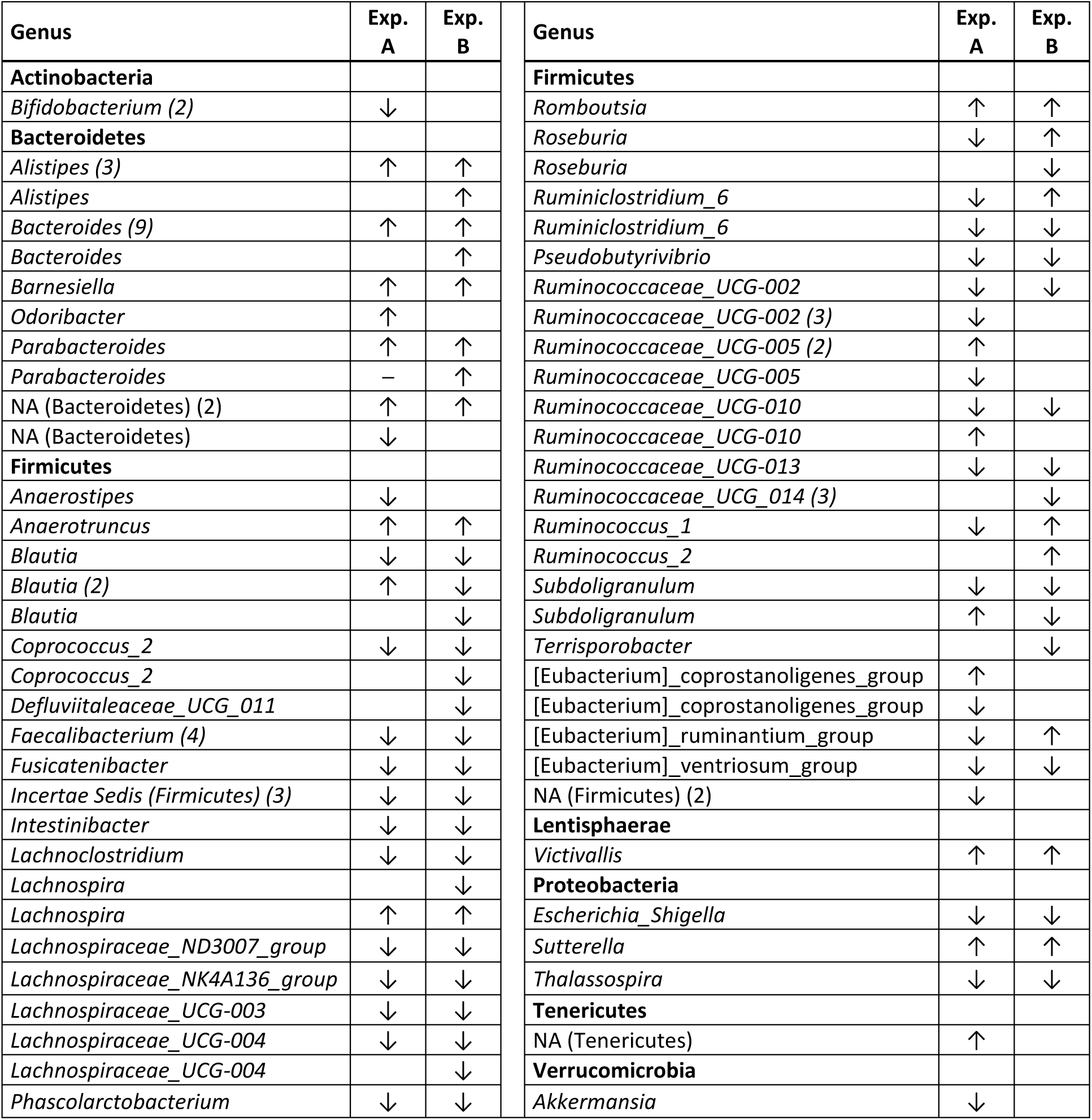
Summary of significantly changed RSVs in experiments A and B (100 most abundant RSVs; LEfSe). RSVs which increased (↑) or decreased (↓) in relative abundance (concordant results for all concentrations) with PMA treatment are listed. Number of RSVs for each genus are given in brackets if there were more than one. One RSV which was present (*Parabacteroides*) but did not change significantly in with PMA treatment (Experiment A) is marked with –. RSVs which were not present in one of the experiments are empty. Numbers after genus information depicts the number of RSVs giving similar result. Full information on the 100 most abundant identified RSVs and results including PMA concentrations is available in Supplementary Table 1 and Supplementary Table 2.

Notably, in both experiments we identified four proteobacterial RSVs (*Bilophila*, *Escherichia/Shigella, Sutterella,* and *Thalassospira)*, and in both experiments *Escherichia/Shigella* and *Thalassospira* were relatively decreased and *Sutterella* was increased with PMA treatment. In both experiments, the abundance of genus *Bilophila* did not alter. These results indicate that *Escherichia/Shigella* and *Thalassospira* are found in gut also in nonviable state and their abundance can be overemphasized.

### Concentration of 100 μM PMA is sufficient to discriminate intact and dead cells in stool samples

We were interested in the optimal concentration of PMA needed to mask the dead cells in stool samples, and studied with which concentration the significant changes in relative abundance are visible at RSV level. Out of 100 most abundant RSVs in experiment A, 75 RSVs changed significantly with PMA treatment; the change occurred only in 6 RSVs with 10 μM PMA concentration. In 14 RSVs (19%) the change was significant already with 20 μM PMA, 30 RSVs with 30 μM PMA, 38 RSVs with 40 μM PMA, 26 RSVs with 50 μM PMA, 45 RSVs with 60 μM PMA, 43 RSVs with 70 μM PMA, 31 RSVs with 80 μM PMA, 45 with RSVs 90 μM PMA, 52 RSVs 100 μM, suggesting that in this experiment, 100 μM of tested concentrations was most efficient in masking the nonviable cells. A graphical overview on experiment A is given in Figure 4.

**Figure 4.**
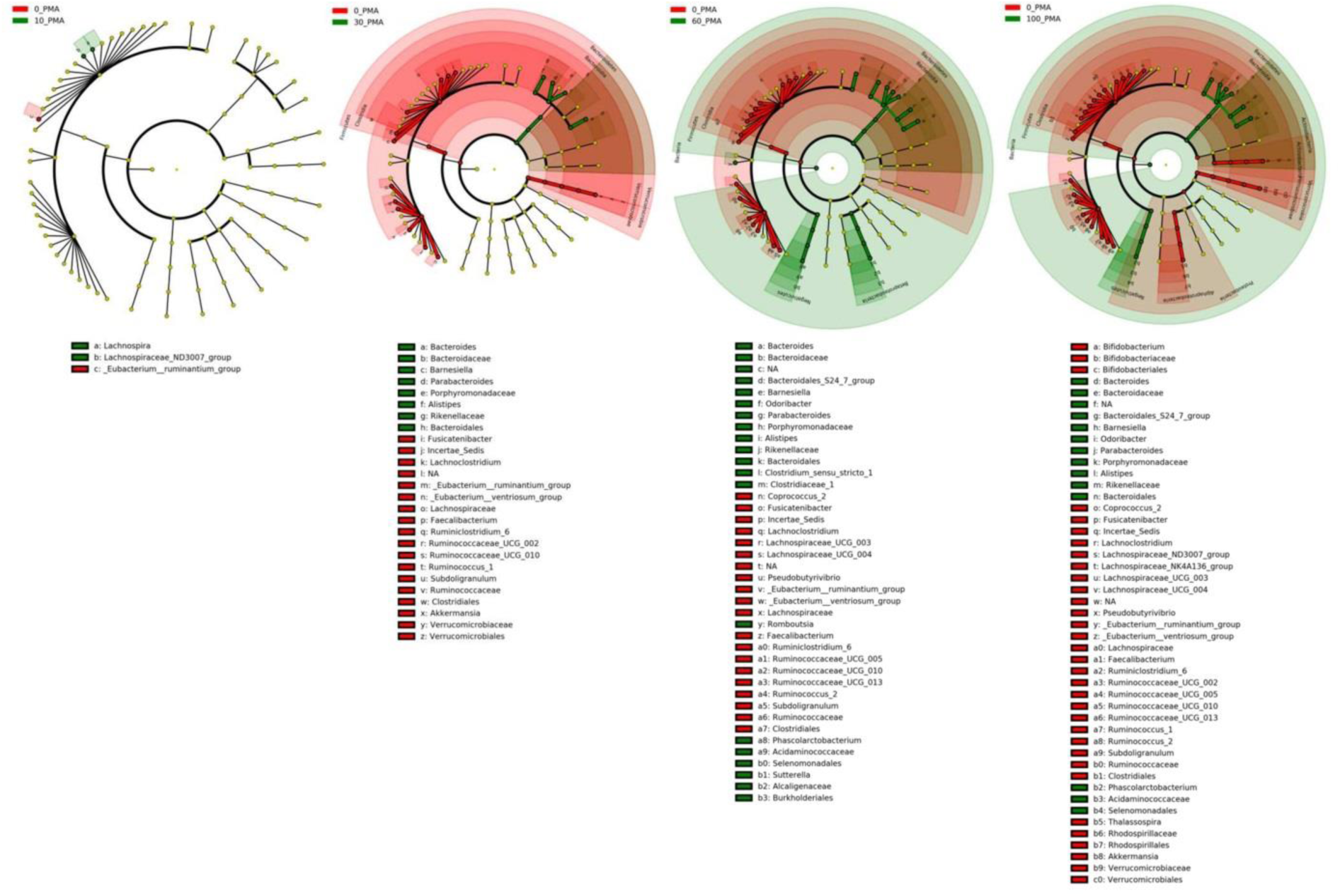
Graphical summary of significantly changed RSVs (100 most abundant, LefSE analyses) in experiment A at concentrations of 10, 30, 60 and 100 µM PMA. RSVs which increased or decreased in relative abundance after PMA treatment are given in green and red, respectively. Full information on the 100 most relatively abundant identified RSVs and results including PMA concentrations is available in Supplementary Table 1.

In experiment B we tested if increasing the amount of PMA concentration up to 300 μM would significantly change the community membership at RSV level. Of 100 most abundant RSVs which were analysed, 68 RSVs changed significantly with PMA treatment. We applied PMA at four different concentrations (50 μM, 100 μM, 150 μM, and 300 μM), and found that 49 RSVs at 50 μM PMA concentration, 44 RSVs at 100 μM PMA concentration, 43 RSVs at 150 μM PMA concentration, and 42 RSVs at 300 μM PMA concentration were increased or decreased significantly. Interestingly, for 59 out of 68 RSVs (87%) the significant change occurred already at 50 or 100 μM PMA, suggesting that using a larger concentration does not substantially affect the community profile on RSV level.

### Firmicutes/Bacteroidetes ratio decreased significantly upon PMA treatment

As detected with LEfSe analysis most of the RSVs belonging to phylum Firmicutes decreased in relative abundance with PMA treatment, and RSVs in phylum Bacteroidetes increased. As Firmicutes/Bacteroidetes ratio in human gut microbiota has been connected to human health and disease, we pursued to look deeper into how the PMA treatment changed this ratio. We found that the Firmicutes/Bacteroidetes ratio decreased by 21% in experiment A at a concentration of 30 μM PMA (Student’s t-test p=0.031), and even 31% at 100 μM of PMA (Student’s t-test p=0.018). In experiment B the Firmicutes/Bacteroidetes ratio did not further decrease with increasing PMA concentration, but a statistically significant 30-33% reduction compared to 0 μM PMA was observed with all PMA concentrations (t-test p values range 0.0004 – 0.04) (Figure 5.).

**Figure 5.**
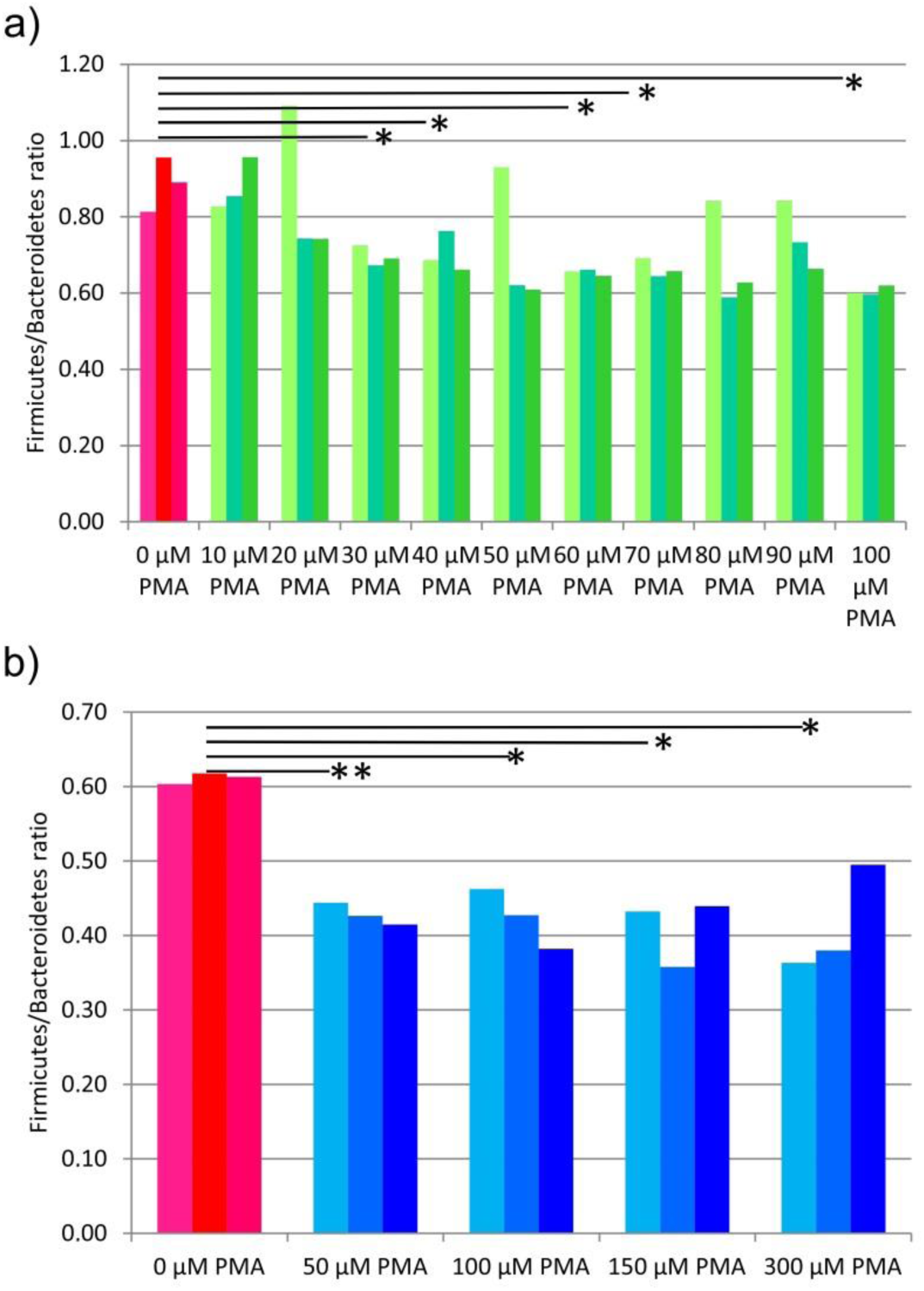
Firmicutes/Bacteroidetes ratio decreased significantly with PMA treatment. a) Experiment A, b) experiment B. The stars above the bars depict the statistical significance of results: when p-value is <0.05, it is shown with *, and when p-value is <0.01, it is shown with **.

## Discussion

The purpose of this study was to analyze the correlation of PMA concentration and the microbial composition after NGS in complex biosamples such as stool, with the goal to establish a guideline for future applications.

Most studies using PMA with stool samples were performed with standard PMA concentration of 50 μM. However, testing the effect of increasing PMA concentrations on stool samples, and determining the relative abundances in correlation with live / dead status of specific microorganisms has not been studied before. Here, we applied Illumina MiSeq sequencing to decipher the effect of PMA treatment on turbid stool samples and their diverse microbial community.

This study revealed that PMA treatment significantly decreases microbial richness and diversity (alpha diversity) and alters the microbial community profile (beta diversity). The magnitude of these effects was found to be concentration dependent. Furthermore, the increase and decrease in relative abundance after PMA treatment was taxon-specific, revealing a significantly different composition of the microbial community when only intact cells were targeted.

A decrease in diversity suggests that either certain taxa totally disappeared (richness), or the evenness decreased when the relative abundances of different taxa shifted upon PMA treatment. In our experiments, the majority of 100 most abundant RSVs either increased or decreased statistically significantly in relative abundance at one or more applied PMA concentrations, but only few RSVs completely disappeared. This suggests that PMA treatment particularly affected the evenness of the microbial community. Furthermore, the change in beta diversity suggests that the community profile shifted upon PMA treatment: certain taxonomic groups became more abundant, some less abundant, and some were not detectable any more. This indicates that they were not intact, but represented in the non PMA treated samples only by fragments of free DNA or DNA within cells with fractured cell walls. Thus, we state that without using any live/dead distinction method, the active and significant microbial diversity in stool might be miscalculated.

The effect of PMA on both alpha and beta diversity was clearly concentration dependent. In particular, the application higher than 100μM PMA showed a significant decrease in the alpha diversity compared to untreated samples. Additionally, the community profile shift was stronger with higher PMA concentrations, suggesting that low PMA concentrations only can mask part of the free DNA or DNA inside fractured cells, because of turbidity.

We observed that the increase and decrease in relative abundance after PMA treatment was taxon specific, revealing a significantly different composition of the microbial community when only intact cells were targeted. Particularly the relative abundance of Firmicutes was significantly reduced and the relative abundance of Bacteroidetes significantly increased when PMA treatment was applied, leading to the conclusion that without any live/dead staining or viability testing the number of Firmicutes might be significantly overestimated while Bacteroidetes are underestimated. In particular, the Firmicutes/Bacteroidetes ratio has been connected to human health in a number of publications ^34-36^.

Gram-positive Firmicutes are predominantly associated with energy harvest from diet^37, 38^, while Gram-negative Bacteroidetes are known to perform several beneficial processes related to degradation of complex sugars and proteins into short chain fatty acids (SCFAs)^39^, and the relative proportion of Bacteroidetes has been observed to be decreased in obesity ^34^. In the Bacteroidetes phylum, we detected a systematic and significant increase of the genera *Bacteroides, Alistipes,* and *Parabacteroides* upon PMA treatment. *Bacteroides* comprises the most substantial portion of the human gastrointestinal microbial community, and is known to play a fundamental role in processing complex molecules to succinate and acetate (major fermentation products) ^40^ in the host intestine ^41, 42^, and it contributes to the lean phenotype ^43^. *Alistipes,* and *Parabacteroides* used to belong to genus *Bacteroides* and thus share many metabolic features, including fermentation of complex sugars, and production of acetic and succinic acids^44-46^.

Within the phylum Firmicutes we observed a systematic decrease of the genus *Faecalibacterium*, which is one of the most abundant and important commensal bacteria of the human gut microbiota. *Faecalibacterium* is known to produce butyrate and other short-chain fatty acids through the fermentation of dietary fiber and to boost the immune system. Additionally, relatively low levels of *F. prausnitzii* in human intestines were associated with e.g. inflammatory bowel diseases (Crohn’s disease) ^47^ and ulcerative colitis ^48^, obesity ^49^, asthma ^50^ and depression ^51^.

Besides *Faecalibacterium*, also other important human gut bacteria were significantly affected by the PMA treatment: the genera *Bifidobacterium* and *Akkermansia* both decreased in relative abundance upon PMA treatment. These bacteria are known to carry beneficial properties for human health and their application as probiotics has been extensively studied. For example *Bifidobacterium* has been connected to beneficial psychological effects ^52^ better mental health ^53^, and allergy prevention ^54^. It is suggested to be effective at protecting against infectious diseases ^55^, whereas *Akkermansia* is known to stimulate immune system, improve gut barrier function ^56-58^, and prevent the development of obesity (fat mass development, insulin resistance and dyslipidemia) in mice studies ^58^. Additionally, the level of *Akkermansia muciniphila* was reported to be drastically reduced in patients with Crohn’s disease and ulcerative colitis, and suggested to be a health biomarker ^59^.

Based on our study, supported by the results of a study aiming at the discrimination of viable and dead fecal Bacteroidales by quantitative PCR ^60^, we suggest using a concentration of 100 μM of PMA to efficiently block background DNA from open or broken cells in fecal samples.

Although human faeces samples do not reflect the situation at the respective site within the intestine ^61, 62^, they are used for numerous microbiome analyses and correlation studies of microbial composition and health or disease. Excluding the background signals by PMA treatment can increase the impact of such studies, as the intact microorganisms might mirror the *in situ* situation in more detail.

### Limitations of the study

The stool samples were immediately processed and handled in the dark throughout the experiments. However, the samples were exposed to air (21% oxygen) for a short period of time during dilution and PMA treatment, which might affect the viability of the microbial community ^2^. However, we cannot think of any chemical process, which would, in the dark but in presence of air, induce immediate cell wall/cell membrane damage, and thus affect the outcome of the PMA treatment ^63^, so that we did not evaluate the impact of the oxygen in the ambient air on our results. In this regard, the study setup was performed according to feasibility and common practice, and the samples were treated with PMA without delay, but under normal atmosphere.

## Conclusion

Treatment with propidium monoazide is a relatively simple and cheap protocol to exclude background DNA from damaged cells in DNA-based microbiome studies. Herein, we propose to use 100 μM PMA for optimal live/dead distinction in stool microbiome analyses, as we identified the standard concentration of 50 μM to be less effective in such turbid samples. In our study, we found that the intact, and thus most likely viable fraction of the microbial community, differs significantly from the overall microbial profile. This finding indicates, that many previous studies might have over- or underestimated the importance of key microbial species in gut samples, as they did not consider the live/dead situation of the cells.

## Declarations

### Ethics approval and consent to participate

Research involving human material was performed in accordance with the Declaration of Helsinki and was approved by the local ethics committees (the Ethics Committee at the Medical University of Graz, Graz, Austria). Stool samples have been obtained covered by the ethics vote 27-151 ex 14/15.

### Consent for publication

Not applicable.

### Availability of data and material

Sequence data were submitted to the European Nucleotide Archive (ENA) with the study accession number PRJEB25855.

## Competing interests

The authors declare that the research was conducted in the absence of any commercial or financial relationships that could be construed as a potential conflict of interest.

## Funding

This study was supported by the Kulturamt Stadt Graz (“Life-dead analyses with stool microbiomes”) and BioTechMed-Graz. MM was trained within the frame of the Ph.D. Program Molecular Medicine of the Medical University of Graz.

## Authors’ Contributions

AP, MM, CME conceived and designed the study. AP, MM, MB, LW collected and processed the samples, KK, AP, CME analysed and interpreted the data, and AKP, KK, CME drafted and revised the manuscript. All authors approved the final version of the manuscript.

## Acknowledgements

We thank the City of Graz cultural office, BioTechMed-Graz, and Ph.D. Program Molecular Medicine of the Medical University of Graz for funding.

## Additional files

**Additional tables**

### Additional Table 1

Microsoft Excel File Format, **.xlsx**

**Description of data**: Summary of 100most abundant taxa in Experiment A, and the changes in relative abundance upon PMA treatment. Bacterial taxa (RSVs) that increased are marked in green, and taxa that decreased are marked in red.

### Additional Table 2

Microsoft Excel File Format, **.xlsx**

**Description of data**: Summary of 100most abundant taxa in Experiment B, and the changes in relative abundance upon PMA treatment. Bacterial taxa (RSVs) that increased are marked in green, and taxa that decreased are marked in red.

